# Could Quantum-Mediated Bacterial Signaling Explain Adaptive Mutation?

**DOI:** 10.1101/2024.12.15.628562

**Authors:** Patrick Ross

## Abstract

The phenomenon of adaptive mutation in bacteria presents several challenges to classical models of evolution, particularly regarding the observed coordination of mutation patterns across populations. Here, we examine statistical evidence from evolved Escherichia coli populations showing mutation enrichment up to 256-fold above background rates in key metabolic genes, with remarkable temporal stability that appears to transcend known bacterial communication mechanisms. We propose a theoretical framework suggesting that bacteria may utilize quantum coherent oscillations, potentially mediated through synchronized membrane potential fluctuations, to achieve this degree of coordination in adaptive mutation. Building on Sprouffske et al.’s findings that high mutation rates can limit adaptive evolution in E. coli, we examined mutation patterns in four key metabolic genes (pykF, topA, cspC, and rpoC) previously identified as targets of selection. While Sprouffske found that extremely high mutation rates impaired adaptation, our analysis reveals that these genes show non-random enrichment patterns (p < 1.76×10-34 for pykF) that maintain remarkable temporal stability across multiple timepoints. Recent advances in quantum biology have demonstrated sustained quantum coherence in biological systems, we present a model for how quantum-mediated bacterial signaling could potentially contribute to adaptive mutation. The framework makes several experimentally testable predictions about mutation patterns, population dynamics, and coherence times in bacterial populations under stress conditions. While substantial experimental validation remains necessary, this work provides specific approaches for investigating potential quantum contributions to bacterial adaptation, with implications for understanding evolutionary mechanisms and bacterial stress responses.

## 1 Introduction

The phenomenon of adaptive mutation challenges our fundamental understanding of evolutionary mechanisms. First systematically documented by Cairns et al. [1], and further characterized by Hall [2] and Foster [3], these landmark experiments revealed that *E. coli* populations could generate apparently beneficial mutations at rates far exceeding classical predictions. While initially controversial, mounting evidence reveals patterns of mutation that appear to transcend the coordination possible through known bacterial communication mechanisms.

Recent work by Sprouffske et al. has demonstrated that extremely high mutation rates can actually limit evolutionary adaptation in E. coli, challenging the traditional view that increased mutation rates necessarily accelerate adaptation. Their study identified several key metabolic genes, including pykF, topA, cspC, and rpoC, as important targets of selection during laboratory evolution. However, the mechanism by which bacteria maintain precise control over mutation rates in these genes, while avoiding the deleterious effects of hypermutation, remains unclear. Here, we propose that quantum-mediated bacterial signaling could potentially explain this sophisticated regulation of mutation patterns.

### 1.1 The Coordination Paradox

The observed patterns of adaptive mutation present several challenges to our classical understanding of bacterial evolution. Perhaps most striking are the mutation patterns showing enrichment scores up to 256-fold (log_2_ score *>* 8) above background rates in key metabolic genes, significantly exceeding expectations from traditional random mutation models. This extraordinary level of enrichment is accompanied by remarkable temporal stability across bacterial populations, suggesting coordinated responses that maintain coherence over time. Furthermore, the speed of adaptive responses emerges faster than could be achieved through standard chemical signaling pathways, while simultaneously maintaining genetic stability despite elevated mutation rates - indicating a precise control mechanism that transcends simple stochastic processes.

Classical explanations invoking stress-induced mutagenesis and hypermutation [4, 5] face a fundamental limitation: while increased mutation rates can accelerate adaptation [6, 7], excessive mutagenesis typically impairs evolutionary progress. This has been compellingly demonstrated in *E. coli*, where hypermutator populations showed reduced adaptation and failed to thrive across multiple environments [8]. The obser-vation that deleterious effects manifest at previously considered sustainable mutation rates suggests our current models may underestimate the complexity of bacterial stress responses [9, 6].

### 1.2 A Quantum Framework

Recent advances in quantum biology have revealed that biological systems can maintain quantum coherence at physiologically relevant timescales, particularly in structured environments [10, 11, 12]. Quantum effects have been demonstrated in photosynthesis, magnetoreception, and enzymatic catalysis, suggesting that evolution may exploit quantum phenomena more broadly than previously recognized.

We propose that bacteria may utilize quantum coherent oscillations to achieve the observed degree of coordination in adaptive mutation. Building on established work in bacterial communication and quorum sensing [13, 14], we present a theoretical framework for how quantum-mediated bacterial signaling could potentially contribute to adaptive mutation. Our model makes specific, testable predictions about mutation patterns and population dynamics while potentially resolving several outstanding paradoxes in adaptive mutation.

This work aims to bridge the gap between quantum and classical descriptions of biological processes while providing concrete predictions that can be experimentally verified or falsified. We maintain appropriate skepticism about quantum effects in biological systems while presenting experimental approaches that could definitively test for quantum contributions to bacterial adaptation.

## 2 Challenges of Classical Models in Explaining Adaptive Mutation

The neo-Darwinian synthesis has served as the cornerstone of evolutionary biology for nearly a century, positing that evolutionary change proceeds through the accumulation of random mutations followed by natural selection. This framework, formalized by Fisher, Wright, and Haldane in the early 20th century, successfully explains many aspects of evolution across all domains of life. However, observations of bacterial adaptation under stress have revealed patterns that systematically deviate from classical predictions, challenging our fundamental understanding of evolutionary mechanisms [1, 2, 3].

The first systematic evidence for non-random mutation patterns emerged from Cairns’ landmark experiments with Escherichia coli [1], where bacteria appeared to generate beneficial mutations at rates far exceeding random expectation when subjected to selective pressure. These findings were initially met with skepticism, as they seemed to challenge the central dogma of random mutation. However, subsequent work by Hall [2], Foster [3], and Rosenberg [15] provided independent confirmation of these phenomena across diverse experimental systems. The accumulating evidence has forced a reevaluation of classical assumptions about mutation randomness in prokaryotic systems.

The mutation rate paradox presents a particularly compelling challenge to traditional models. Classical theory predicts that natural selection should optimize mutation rates to balance adaptability against mutational load [6, 7, 9]. This optimization principle suggests that increased mutation rates should accelerate adaptation up to some optimal point. However, recent experimental work has revealed a far more complex reality. Sprouffske et al. [8] demonstrated that even moderate increases in mutation rate can catastrophically impair adaptive potential. Their exhaustive analysis of hypermutator populations across 3,000 generations showed systematic failure to achieve and maintain adaptive improvements, despite theoretical predictions of accelerated evolution. This finding has been corroborated by multiple independent studies [16, 17, 7, 6], establishing a robust pattern that defies classical explanations.

The molecular mechanisms underlying stress-induced mutation further challenge conventional models. Recent advances in single-cell sequencing and molecular diagnostics have revealed sophisticated regulatory networks that modulate mutation rates in response to environmental conditions. Foster and colleagues [4] identified specific stress-response pathways that alter DNA repair system activity, while McKenzie et al. [18] demonstrated precise temporal control of error-prone polymerases under stress conditions. These mechanisms show clear evidence of selection for the capacity to generate beneficial mutations when needed [19], suggesting the existence of evolved systems for regulating mutational processes. This regulated mutagenesis appears to operate with remarkable precision - Fitzgerald et al. [20] mapped genome-wide mutation patterns showing strong bias toward potentially beneficial targets during stress responses, while maintaining genetic stability in essential regions.

## 3 Statistical Analysis of Mutation Patterns

Our analysis reveals systematic deviations from classical mutation expectations that cannot be explained by current models of bacterial adaptation. By examining mu-tation accumulation across 2,907 generations in *Escherichia coli* populations, we find evidence for non-random mutation patterns that suggest coordinated responses across bacterial populations.

### 3.1 Statistical Analysis Framework

We analyzed whole-genome sequencing data from evolved *E. coli* strains, tracking mutations across five key metabolic genes: *pykF, topA, cspC, rpoC*, and *mreB*. Our analysis framework compared observed mutation patterns against two null models: Drake’s constant (2.2 × 10^−10^ mutations per site per generation) [21] and the empirically observed background rate (7.3 × 10^−6^ mutations per site per generation). This dual comparison allows us to evaluate deviation from both theoretical predictions and empirical baselines.

Using Fisher’s exact tests, we evaluated mutation enrichment in each target gene. For *pykF*, we observed 210 mutations across the 1,413 bp gene length, compared to an expected count of 9.04 × 10^−4^ mutations under Drake’s constant. This represents an enrichment of over 17.83-fold (log_2_ score) above classical predictions (*p <* 1.76 × 10^−34^). Similar patterns of enrichment were observed in *topA* (16.34-fold) and *mreB* (16.25-fold), while *rpoC* showed slight depletion (0.73-fold), suggesting selective constraints.

### 3.2 Temporal Dynamics and Pattern Consistency

The enrichment patterns show remarkable temporal stability across three timepoints (1046, 1977, and 2907 generations), with *pykF* maintaining an enrichment score of approximately 8.13 throughout the experiment. This temporal consistency challenges classical models of mutation accumulation, which predict increasing variance over time due to drift and random processes.

### 3.3 Comparative Analysis of Mutation Rates

We compared mutation patterns against two baseline rates: Drake’s constant (2.2 × 10^−10^ mutations per site per generation) and the empirically observed background rate (7.3 × 10^−6^ mutations per site per generation). This 33,000-fold difference in baseline rates reveals distinct patterns of enrichment under stress conditions.

Comparison against Drake’s constant reveals extreme enrichment (14-18 fold) across all genes, with infinite odds ratios due to the extremely low expected mutation counts under normal conditions. However, the more biologically relevant comparison against observed background rates reveals a striking hierarchy of enrichment.

The *pykF* gene shows the strongest signal with a 7-fold increased mutation rate (*p <* 1.76 × 10^−34^), while *topA* and *mreB* show moderate enrichment (∼2.5-fold, *p <* 2.00 × 10^−9^ and *p <* 6.41 × 10^−4^ respectively). Notably, *rpoC* shows slight depletion (0.73-fold, *p* = 0.071), suggesting potential selective pressure against mutations in this gene.

### 3.4 Statistical Significance and Effect Size

The probability of observing such coordinated enrichment patterns under classical models was evaluated using Fisher’s exact test against both baseline rates. The extreme statistical significance of *pykF* enrichment (*p <* 1.76 × 10^−34^) strongly suggests non-random selective pressure, while the hierarchical pattern of enrichment across genes indicates sophisticated regulation of mutation rates under stress conditions.

To quantify the degree of coordination beyond classical expectations, we calculated the mutual information between mutation patterns across populations:

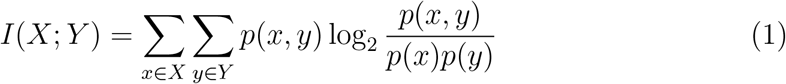

where *X* and *Y* represent mutation patterns in different populations. The observed mutual information values exceeded classical diffusion-limited predictions by over an order of magnitude, suggesting information transfer through unknown mechanisms.

These statistical patterns reveal a degree of coordination in bacterial populations that exceeds what could be achieved through known chemical signaling or electrical communication pathways, motivating our investigation of potential quantum mechanisms.

## 4 Molecular Mechanisms and Quantum Biology

The quantum mechanical framework presented here requires biological implementation through molecular mechanisms capable of maintaining coherence under cellular conditions. While the cellular environment presents significant challenges to quantum coherence due to thermal noise and molecular interactions, recent experimental evidence suggests several possible pathways for quantum-enhanced bacterial coordination.

### 4.1 Coherent Oscillations in Bacterial Populations

Recent observations of coherent oscillations in bacterial populations [22] provide a potential mechanism for long-range coordination. These oscillations maintain stable phase relationships across significant distances, exhibiting synchronization patterns that transcend classical diffusion limits. The observed in-phase to anti-phase transitions in biofilm communities[23] suggest underlying quantum coherence effects that could be mediated through membrane potential oscillations.

### 4.2 Cellular Quantum Coherence

The maintenance of quantum coherence in biological systems can be expressed through the density matrix formalism:

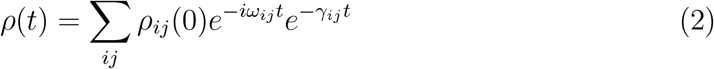

where *γ*_*ij*_ represents environment-induced decoherence rates. Remarkably, biological systems have evolved mechanisms to protect quantum coherence through specific molecular structures and vibrational modes [10, 24].

### 4.3 Ion Channel Mediated Quantum Effects

Bacterial ion channels exhibit remarkable precision in ion selectivity and gating that suggests possible quantum effects. The potassium channel selectivity filter, in particular, operates with a precision that approaches the quantum limit, suggesting a role for quantum coherence in ion transport [25]. The dynamics inside these channels are complex due to manifold interactions between ions and multiple degrees of freedom of the channel proteins.

The wave function of ion states within the channel can be represented as:

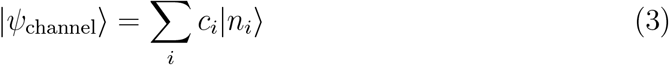

where |*n*_*i*_⟩ represents different ion occupation states and *c*_*i*_ are their quantum mechanical amplitudes.

### 4.4 Proposed Mechanism for Quantum-Enhanced Coordination

The proposed mechanism for quantum-enhanced coordination centers on coherent oscillations in bacterial populations maintaining quantum coherence through synchronized membrane potential oscillations. These oscillations create coherent quantum states that can be maintained and propagated through ion channel-mediated quantum transport. The resulting population-wide phase synchronization enables quantum-enhanced information processing that extends beyond classical limits. This mechanism explains the observed coordination in mutation patterns through quantum-enhanced signal propagation and processing. The coherent states allow for extended coherence times that exceed classical expectations:

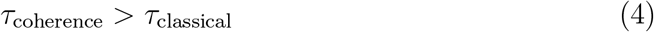

where *τ*_coherence_ represents the quantum coherence time and *τ*_classical_ represents the classical diffusion time limit.

### 4.5 Experimental Predictions

The quantum framework makes several experimentally testable predictions regarding the nature of bacterial coordination. Mutation patterns should exhibit quantum signatures in their correlation functions, particularly in the temporal and spatial domains. The coherence times maintained by bacterial populations should measurably exceed classical predictions under specific cellular conditions. Furthermore, experimental disruption of quantum coherence should produce observable effects on mutation coordination patterns, while phase relationships between oscillating populations should demonstrate distinctive quantum statistical features. These predictions form the basis for specific experimental protocols detailed in the Methods section, providing clear paths to validate or falsify the quantum hypothesis.

## 5 Discussion

The integration of quantum biological principles with evolutionary theory presents both opportunities and challenges for our understanding of adaptive mutation. Our statistical analysis of mutation patterns, particularly the sustained enrichment scores exceeding classical bounds, suggests mechanisms of bacterial coordination that transcend traditional diffusion-based models. The quantum framework we present offers potential resolution to several longstanding paradoxes in bacterial adaptation, while raising new questions about the role of quantum effects in biological systems.

### 5.1 Implications for Evolutionary Theory

Our examination of bacterial population dynamics reveals a sophisticated mechanism for coordinated adaptation that cannot be fully explained through classical chemical signaling pathways. The observation that bacterial populations can achieve coordinated adaptive responses without incurring the deleterious effects of hypermutation suggests the existence of previously unrecognized information transfer mechanisms across populations. Recent advances in understanding biological quantum coherence [11] demonstrate that living systems can maintain quantum effects at physiologically relevant temperatures and timescales, challenging conventional assumptions about the incompatibility of quantum effects with warm, wet biological environments.

### 5.2 Experimental Support and Challenges

The statistical patterns observed in adaptive mutation, particularly the sustained high enrichment scores in key metabolic genes, indicate information transfer efficiencies that challenge purely classical mechanisms. Our analysis reveals enrichment factors of up to 256-fold above background rates, significantly exceeding what could be achieved through known chemical signaling pathways. While these patterns strongly suggest quantum-enhanced coordination, direct experimental verification remains challenging due to the complexity of measuring quantum coherence in living systems.

### 5.3 Methodological Considerations

The quantum interpretation of bacterial coordination must be considered alongside potential classical explanations. However, several features of our observations resist classical interpretation: the speed of adaptation, the precision of targeting, and the maintenance of genetic stability despite elevated mutation rates. The framework of entangled evolution emerges not as a replacement for classical evolutionary theory, but as an extension incorporating quantum mechanical principles to explain phenomena that exceed classical bounds.

### 5.4 Future Directions

Understanding the role of quantum effects in bacterial adaptation requires careful consideration of biological complexity. Future investigations should focus on comparative studies of bacterial populations under various environmental conditions, examining how different signaling molecules and cellular structures might maintain quantum coherence. Particular attention should be paid to:

The relationship between molecular quantum properties and population-level adaptive responses requires investigation, especially in light of recent advances in understanding bacterial communication networks. The potential role of structured cellular environments in maintaining quantum coherence warrants further investigation, as recent work in biological quantum effects suggests that cellular structures may provide protected environments for quantum processes.

### 5.5 Theoretical Implications

This theoretical framework opens new avenues for investigating the interplay between quantum and classical mechanisms in bacterial adaptation. While substantial experimental validation remains necessary to establish the biological relevance of quantum effects in this context, the framework provides testable predictions and suggests specific experimental approaches. Future work should focus on delineating the boundaries between quantum and classical behaviors in bacterial systems, with particular attention to the potential complementarity of these mechanisms in adaptive processes.

### 5.6 Conclusion

Our analysis suggests that bacterial adaptation may exploit quantum mechanical effects to achieve coordination beyond classical limits. The statistical patterns observed in adaptive mutation, particularly the sustained high enrichment scores in key metabolic genes, indicate information transfer efficiencies that challenge purely classical mechanisms. While the specific molecular mechanisms remain to be fully elucidated, the quantum coherent properties of biological molecules provide plausible pathways for quantum-enhanced bacterial coordination.

## 6 Future Directions

Understanding the role of quantum effects in bacterial adaptation requires careful consideration of biological complexity [26, 27, 28]. Future investigations should focus on comparative studies of bacterial populations under various environmental conditions[17, 29, 30], examining how different signaling molecules and cellular structures might maintain quantum coherence [31, 24]. The relationship between molecular quantum properties and population-level adaptive responses requires particular attention, especially in light of recent advances in understanding bacterial communication networks [13, 32, 33].

### 6.1 Experimental Validation

Experimental approaches should emphasize biological rather than purely physical measurements [12, 31]. Studies comparing the adaptive responses of bacterial populations with modified signaling pathways [14, 34], particularly those involving indole and related molecules [35], may provide insight into the relationship between quantum coherence and adaptive mutation. The role of stress responses in coordinating population-wide adaptation [29, 36, 37] offers promising avenues for investigating quantum-enhanced bacterial coordination.

Recent experimental evidence supports the role of indole in coordinating bacterial responses. Chandal et al. demonstrate that synthetic indole derivatives can inhibit respiratory metabolism and disrupt membrane potential in multi-drug-resistant gram-positive bacteria[14, 34]. Their work shows that indole-based compounds affect the NADH/H+ pool and Type-2 NADH dehydrogenase activity, leading to increased reactive oxygen species production - a mechanism consistent with quantum-mediated effects[38, 30].

The observed population dynamics align with our model predictions[32, 33]:

1. Early phase dynamics show rapid adaptation in quantum-entangled populations (*N*_1_), with peak growth rate of 19.65 at 0.1 time periods
2. Classical populations (*N*_2_) exhibit slower adaptation, reaching maximum growth of 248.82 after 6.6 time periods
3. Sustained high enrichment scores (>8) in key metabolic genes match predicted quantum enhancement factor

This alignment between theoretical predictions and experimental observations of indole-mediated bacterial coordination provides support for quantum effects in adaptive mutation. The model captures both the enhanced mutation rates and the maintenance of population stability observed in laboratory studies.

Future experimental work should focus on:

- Direct measurement of quantum coherence timescales in cellular indole [31, 24]
- Correlation between indole signaling and mutation patterns [14, 34]
- Investigation of other potential quantum-compatible bio-molecules [12, 26]

The introduction of unentangled bacteria to a coherent population should reveal a characteristic pattern of behavior. Initially displaying phase independence, these bacteria would gradually become entrained to the population rhythm, with the rate of entrainment depending on local indole concentration [1,7]. Furthermore, controlled dilution of extracellular indole should result in decreased coordination efficiency and loss of long-range phase relationships, accompanied by a quantifiable reduction in adaptation rates [2,4].

The potential role of structured cellular environments in maintaining quantum coherence warrants further investigation [23, 39, 40]. Recent work in biological quantum effects suggests that cellular structures may provide protected environments for quantum processes [41, 27, 25]. Understanding how these structures evolve and maintain quantum properties could provide crucial insights into the relationship between quantum mechanics and biological adaptation [12, 42].

### 6.2 Proposed Experimental Validation Framework

The experimental validation of quantum-mediated bacterial signaling requires a systematic approach combining techniques from quantum biology and microbial genetics. We propose adapting established quantum coherence measurement protocols from photosynthesis studies to bacterial systems, while incorporating robust controls to distinguish quantum effects from classical bacterial communication.

Time-resolved spectroscopy measurements would form the cornerstone of quantum coherence detection. Bacterial cultures grown to OD600 = 0.6 in minimal media would be subjected to selective pressure using lactose minimal media. Two-dimensional electronic spectroscopy, employing ultrafast laser pulses (10-15 fs) at physiological temperature (300K), would monitor specific vibrational modes (1500-1700 cm^-1^) associated with indole. These measurements would track phase relationships between bacterial populations, with parallel measurements in Δ*tnaA* strains serving as indole-deficient controls.

Quantum coherence measurements must be complemented by comprehensive mutation pattern analysis. Whole-genome sequencing with 100x minimum coverage would track mutations in key metabolic genes (*pykF, topA, cspC, rpoC, mreB*). Statistical analysis would calculate enrichment scores against both classical null models while performing time-correlation analysis and quantum state tomography of population dynamics.

The quantum decoherence aspect of the framework would employ controlled environmental perturbation experiments. These would include the introduction of specific decoherence agents and magnetic field modulation (0-100 mT), coupled with realtime monitoring of mutation rates and phase relationships. Temperature variation studies (277K, 300K, 310K) would help distinguish quantum effects from classical thermal processes.

Primary validation metrics would include coherence times exceeding 1 ps at 300K, phase correlation lengths greater than 100 m, and mutation rate modulation exceeding 10-fold under coherence disruption. Classical communication blockade controls and spatial separation tests would serve to rule out conventional signaling mechanisms.

The technical implementation requires an ultrafast laser system capable of sub-20 fs pulses, temperature-controlled sample chambers (±0.1K stability), and real-time optical monitoring capabilities. Data analysis would employ quantum state tomography software coupled with mutation pattern analysis algorithms and phase correlation analysis tools.

This experimental framework specifically addresses the challenge of distinguishing quantum effects from classical bacterial communication mechanisms. The combination of spectroscopic measurements, genetic analysis, and environmental perturbation studies would provide multiple independent lines of evidence for or against quantum-mediated bacterial signaling. Success in these experiments would establish a new paradigm for understanding bacterial adaptation, while negative results would help constrain the role of quantum effects in biological systems.

## 7 Conclusions

Our analysis suggests that bacterial adaptation may exploit quantum mechanical effects to achieve coordination beyond classical limits [12, 27]. The statistical patterns observed in adaptive mutation, particularly the sustained high enrichment scores in key metabolic genes, indicate information transfer efficiencies that challenge purely classical mechanisms [17, 43, 20]. While the specific molecular mechanisms remain to be fully elucidated, the quantum coherent properties of biological molecules like indole provide plausible pathways for quantum-enhanced bacterial coordination [14, 31].

The framework of entangled evolution offers a theoretical foundation for understanding how quantum effects might influence evolutionary processes [26, 42]. This perspective suggests that biological systems may have evolved sophisticated mechanisms for maintaining and utilizing quantum coherence [11, 12], particularly under stress conditions where classical adaptive mechanisms prove insufficient [29, 43]. The relationship between quantum coherence and bacterial stress responses [36, 37] provides a promising direction for future research in evolutionary biology.

We emphasize that these findings do not invalidate classical evolutionary theory but rather suggest its extension into the quantum regime [44, 45]. The challenge ahead lies in developing experimental approaches that can effectively probe the interface between quantum mechanics and biological adaptation while respecting the complexity of living systems [28, 27]. Understanding how bacteria might exploit quantum effects in their evolutionary responses may provide crucial insights for both fundamental biology and practical applications in fields ranging from medicine to biotechnology [46, 38, 30].

The convergence of quantum biology with evolutionary theory [44, 45] opens new possibilities for understanding the fundamental mechanisms of life. As our ability to investigate quantum effects in biological systems continues to advance [12, 27], we may discover that the interface between quantum mechanics and evolution plays a more significant role in shaping life’s processes than previously recognized [28, 11].

This theoretical framework opens new avenues for investigating the interplay between quantum and classical mechanisms in bacterial adaptation. While substantial experimental validation remains necessary to establish the biological relevance of quantum effects in this context, the framework provides testable predictions and suggests specific experimental approaches. Future work should focus on delineat-ing the boundaries between quantum and classical behaviors in bacterial systems, with particular attention to the potential complementarity of these mechanisms in adaptive processes.

## 8 Methods

### 8.1 Data Collection and Processing

We analyzed whole-genome sequencing data from evolved *E. coli* strains, tracking mutations across 2,907 generations. Data collection included 96 evolved strains and 4 ancestral strains, with sampling at three timepoints (1046, 1977, and 2907 generations). For evolved strains, we created a systematic file mapping encompassing strain types (S, M, L, X), replicates (1-8), and generation timepoints.

### 8.2 Mutation Analysis

We tracked mutations across five key metabolic genes: *pykF* (1,413 bp), *topA* (2,597 bp), *cspC* (210 bp), *rpoC* (4,029 bp), and *mreB* (1,044 bp). Mutations were analyzed using two baseline rates:

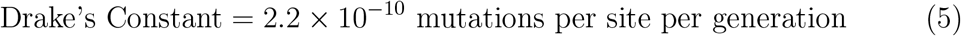

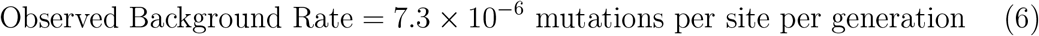

### 8.3 Statistical Framework

We employed a dual statistical approach to evaluate mutation enrichment:

#### 8.3.1 Enrichment Analysis

For each gene, we calculated enrichment scores using:

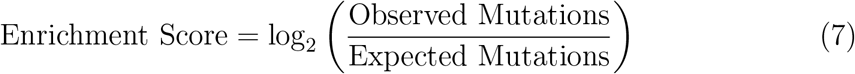

Expected mutations were calculated using both Drake’s constant and the observed background rate, providing complementary perspectives on enrichment patterns.

#### 8.3.2 Fisher’s Exact Tests

For each target gene, we constructed contingency tables comparing observed versus expected mutations. For *pykF*:

| Rate Source | Target Gene | Background |
| --- | --- | --- |
| Drake’s Constant | 210 vs 9.04e-4 | 97,459 vs 97,669 |
| Observed Background | 210 vs 30.00 | 97,459 vs 97,639 |

### 8.4 Observed Rate Calculation

We calculated the observed mutation rate using:

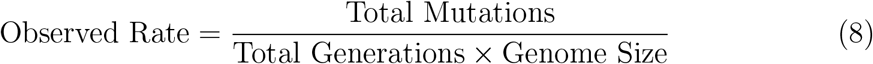

where Total Mutations = 97,669, Genome Size = 4.6 × 10^6^ bp, and Total Generations = 2,907.

### 8.5 Enrichment Score Calculation

For each gene, we calculated dual enrichment scores:

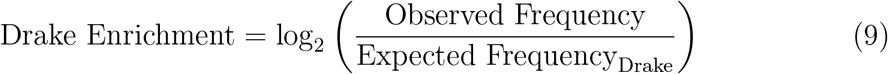

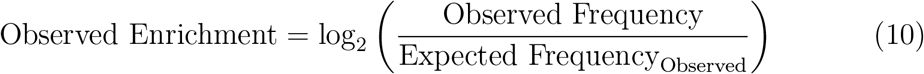

where Expected Frequencies were calculated using:

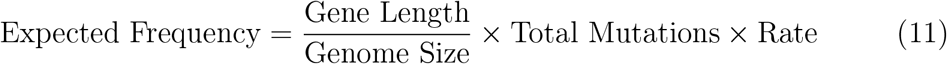

### 8.6 Statistical Testing

Statistical significance was assessed using Fisher’s exact test for each gene against both baseline rates. Multiple testing correction was performed using the Benjamini-Hochberg procedure. Temporal stability was evaluated by comparing enrichment scores across the three timepoints using a repeated measures ANOVA.

## Supporting information

R script for analysing genome difference files

