## Supplementary material for "Could Quantum-Mediated Bacterial Signaling Explain Adaptive Mutation?": R script for analysing genome difference files: mutation_analysis.R

```
# ===============================================
# Setup and Required Packages
# ===============================================
library(dplyr)
library(tidyr)
library(ggplot2)
library(gridExtra)
library(entropy)
library(readr)
library(boot)      # Added for bootstrapping
library(stats)     # Added for additional statistical tests
library(multtest)  # Added for multiple testing correction


# Constants
DRAKES_CONSTANT <- 2.2e-10  # Mutations per site per generation
GENOME_SIZE <- 4.6e6        # E. coli genome size

# ===============================================
# File Processing Functions
# ===============================================
create_file_mapping <- function() {
  # Create evolved strain mapping
  mapping <- data.frame(
    file_num = 1:96,
    prefix = paste0(1:96, "-annot.gd")
  ) %>%
    mutate(
      strain = rep(rep(c("S", "M", "L", "X"), each = 8), 3),
      replicate = rep(1:8, 12),
      generation = rep(c(1046, 1977, 2907), each = 32)
    )
  
  # Add ancestor files
  ancestors <- data.frame(
    file_num = 97:100,
    prefix = c("97_ATGTCA-annot.gd", "97_CCGTCC-annot.gd", "99-annot.gd", "100-annot.gd"),
    strain = c("S", "M", "L", "X"),
    replicate = 0,
    generation = 0
  )
  
  rbind(mapping, ancestors)
}

parse_target_mutations <- function(file_path, target_genes) {
  if (!file.exists(file_path)) {
    warning(paste("File not found:", file_path))
    return(NULL)
  }
  
  lines <- readLines(file_path)
  
  mutations <- data.frame(
    gene_name = character(),
    mutation_type = character(),
    position = numeric(),
    frequency = numeric(),
    stringsAsFactors = FALSE
  )
  
  for (line in lines) {
    if (grepl("^#|^$", line)) next
    if (!any(sapply(target_genes, function(x) grepl(x, line, ignore.case = TRUE)))) next
    
    fields <- strsplit(line, "\t")[[1]]
    mutation_type <- fields[1]
    
    # Extract gene name
    gene_name <- NA
    gene_match <- regexpr("gene_name=([^\\s]+)", line)
    if (gene_match > 0) {
      full_gene_name <- sub("gene_name=", "", regmatches(line, gene_match))
      all_genes <- strsplit(full_gene_name, "/|,")[[1]]
      all_genes <- sub("\\s.*", "", all_genes)
      
      for (gene in target_genes) {
        if (any(grepl(paste0("^", gene, "$"), all_genes))) {
          gene_name <- gene
          break
        }
      }
    }
    
    # Extract position and frequency
    position <- NA
    pos_match <- regexpr("(position|aa_position)=([0-9]+)", line)
    if (pos_match > 0) {
      position <- as.numeric(sub(".*?=", "", regmatches(line, pos_match)))
    }
    
    frequency <- NA
    freq_match <- regexpr("frequency=([0-9.]+)", line)
    if (freq_match > 0) {
      frequency <- as.numeric(sub("frequency=", "", regmatches(line, freq_match)))
    }
    
    if (!is.na(gene_name)) {
      mutations <- rbind(mutations, data.frame(
        gene_name = gene_name,
        mutation_type = mutation_type,
        position = position,
        frequency = frequency,
        stringsAsFactors = FALSE
      ))
    }
  }
  
  return(mutations)
}

# ===============================================
# Analysis Functions
# ===============================================

calculate_observed_rate <- function(mutations_df) {
  total_mutations <- nrow(mutations_df)
  total_generations <- max(mutations_df$generation)
  total_sites <- GENOME_SIZE
  
  return(total_mutations / (total_generations * total_sites))
}


calculate_dual_enrichment <- function(observed_freq, gene_length, observed_rate) {
  TOTAL_MUTATIONS <- 97669
  
  # Calculate expected frequencies based on total mutations
  expected_freq_drake <- (gene_length / GENOME_SIZE) * TOTAL_MUTATIONS
  expected_freq_observed <- (gene_length / GENOME_SIZE) * TOTAL_MUTATIONS * observed_rate
  
  return(list(
    drake_enrichment = log2(observed_freq / expected_freq_drake),
    observed_enrichment = log2(observed_freq / expected_freq_observed)
  ))
}

analyze_mutations <- function(directory_path) {
  target_genes <- c("pykF", "topA", "cspC", "rpoC", "mreB")
  file_mapping <- create_file_mapping()
  all_results <- list()
  background_mutations <- list()  # New list for background mutation data
  
  for (i in 1:nrow(file_mapping)) {
    file_path <- file.path(directory_path, file_mapping$prefix[i])
    
    # Parse target gene mutations
    mutations <- parse_target_mutations(file_path, target_genes)
    
    # Parse all mutations for background comparison
    all_mutations <- parse_all_mutations(file_path)  # New function needed
    
    if (!is.null(mutations) && nrow(mutations) > 0) {
      mutations$file_num <- file_mapping$file_num[i]
      mutations$strain <- file_mapping$strain[i]
      mutations$replicate <- file_mapping$replicate[i]
      mutations$generation <- file_mapping$generation[i]
      all_results[[length(all_results) + 1]] <- mutations
    }
    
    if (!is.null(all_mutations) && nrow(all_mutations) > 0) {
      all_mutations$file_num <- file_mapping$file_num[i]
      all_mutations$strain <- file_mapping$strain[i]
      all_mutations$replicate <- file_mapping$replicate[i]
      all_mutations$generation <- file_mapping$generation[i]
      background_mutations[[length(background_mutations) + 1]] <- all_mutations
    }
  }
  
  if (length(all_results) > 0 && length(background_mutations) > 0) {
    combined_results <- do.call(rbind, all_results)
    combined_background <- do.call(rbind, background_mutations)
    return(list(
      raw_data = combined_results,
      background_data = combined_background
    ))
  }
  return(NULL)
}

analyze_gene_selection <- function(results) {
  mutations <- results$raw_data
  background <- results$background_data
  
  
  # Add this line to calculate observed rate
  observed_rate <- calculate_observed_rate(mutations)
  
  # Calculate counts and probabilities for target genes
  gene_counts <- table(mutations$gene_name, mutations$generation)
  gene_probs <- prop.table(gene_counts, margin = 2)
  
  # Calculate background mutation rates
  background_counts <- table(background$gene_name, background$generation)
  background_probs <- prop.table(background_counts, margin = 2)
  
  # Calculate enrichment scores
  enrichment_scores_drake <- matrix(0, nrow = nrow(gene_probs), ncol = ncol(gene_probs))
  enrichment_scores_observed <- matrix(0, nrow = nrow(gene_probs), ncol = ncol(gene_probs))
  rownames(enrichment_scores_drake) <- rownames(gene_probs)
  colnames(enrichment_scores_drake) <- colnames(gene_probs)
  rownames(enrichment_scores_observed) <- rownames(gene_probs)
  colnames(enrichment_scores_observed) <- colnames(gene_probs)
  
  for(i in 1:nrow(gene_probs)) {
    for(j in 1:ncol(gene_probs)) {
      gene <- rownames(gene_probs)[i]
      gene_len <- gene_lengths$length[gene_lengths$gene_name == gene]
      enrichments <- calculate_dual_enrichment(gene_probs[i,j], gene_len, observed_rate)
      enrichment_scores_drake[i,j] <- enrichments$drake_enrichment
      enrichment_scores_observed[i,j] <- enrichments$observed_enrichment
    }
  }
  
  
  # Print summaries
  print("Mutation counts by gene and generation:")
  print(gene_counts)
  print("\nProbabilities of gene selection by generation:")
  print(gene_probs)
  print("\nEnrichment scores using Drake's constant:")
  print(enrichment_scores_drake)
  print("\nEnrichment scores using observed rate:")
  print(enrichment_scores_observed)
  

  # Add chi-square test for each generation
  chi_sq_results <- list()
  generations <- unique(mutations$generation)
  
  
  return(list(
    counts = gene_counts,
    probabilities = gene_probs,
    enrichment_scores_drake = enrichment_scores_drake,
    enrichment_scores_observed = enrichment_scores_observed,
    observed_mutation_rate = observed_rate,
    chi_square_results = chi_sq_results
  ))
}

# Helper function to parse all mutations (needs to be implemented)
parse_all_mutations <- function(file_path) {
  if (!file.exists(file_path)) {
    warning(paste("File not found:", file_path))
    return(NULL)
  }
  
  lines <- readLines(file_path)
  
  mutations <- data.frame(
    gene_name = character(),
    mutation_type = character(),
    position = numeric(),
    frequency = numeric(),
    stringsAsFactors = FALSE
  )
  
  for (line in lines) {
    if (grepl("^#|^$", line)) next
    
    fields <- strsplit(line, "\t")[[1]]
    mutation_type <- fields[1]
    
    # Extract gene name
    gene_name <- NA
    gene_match <- regexpr("gene_name=([^\\s]+)", line)
    if (gene_match > 0) {
      gene_name <- sub("gene_name=", "", regmatches(line, gene_match))
      gene_name <- sub("\\s.*", "", gene_name)
    }
    
    # Extract position and frequency
    position <- NA
    pos_match <- regexpr("(position|aa_position)=([0-9]+)", line)
    if (pos_match > 0) {
      position <- as.numeric(sub(".*?=", "", regmatches(line, pos_match)))
    }
    
    frequency <- NA
    freq_match <- regexpr("frequency=([0-9.]+)", line)
    if (freq_match > 0) {
      frequency <- as.numeric(sub("frequency=", "", regmatches(line, freq_match)))
    }
    
    if (!is.na(gene_name)) {
      mutations <- rbind(mutations, data.frame(
        gene_name = gene_name,
        mutation_type = mutation_type,
        position = position,
        frequency = frequency,
        stringsAsFactors = FALSE
      ))
    }
  }
  
  return(mutations)
}

# ===============================================
# Quantum-Classical Analysis Integration
# ===============================================

# Gene length data for statistical testing
gene_lengths <- data.frame(
  gene_name = c("pykF", "topA", "cspC", "rpoC", "mreB"),
  length = c(1413, 2597, 210, 4029, 1044)  # lengths in bp
)

perform_fisher_test <- function(mutations, gene_lengths, background_rate) {
  
  # Get observed counts
  observed_counts <- table(mutations$gene_name)
  
  # Get total mutations and generations
  total_mutations <- 97669  # This is our actual total mutations observed
  total_generations <- max(mutations$generation)
  
  # Calculate expected counts based on background rate
  expected_counts <- numeric(length(observed_counts))
  names(expected_counts) <- names(observed_counts)
  
  for (gene in names(observed_counts)) {
    gene_len <- gene_lengths$length[gene_lengths$gene_name == gene]
    # Expected mutations based on background rate
    expected_counts[gene] <- background_rate * gene_len * total_generations
  }
  
  # Perform Fisher's test for each gene
  fisher_results <- list()
  
  for (gene in names(observed_counts)) {
    # Create contingency table comparing to background expectation
    contingency_table <- matrix(
      c(observed_counts[gene], 
        total_mutations - observed_counts[gene],  # Compare to ALL other mutations
        expected_counts[gene], 
        total_mutations - expected_counts[gene]), # Total minus expected in gene
      nrow = 2,
      byrow = TRUE
    )
    
    fisher_test <- fisher.test(contingency_table)
    
    fisher_results[[gene]] <- list(
      gene_name = gene,
      contingency_table = contingency_table,
      p_value = fisher_test$p.value,
      odds_ratio = fisher_test$estimate,
      enrichment = log2(observed_counts[gene] / expected_counts[gene])
    )
  }
  
  return(fisher_results)
}

# Calculate background rate
calculate_background_rate <- function(background_data) {
  # Get total mutations across all generations
  total_mutations <- nrow(background_data)
  
  # Get maximum generation
  max_gen <- max(background_data$generation)
  
  # Calculate rate per site per generation
  rate <- total_mutations / (max_gen * GENOME_SIZE)
  
  # Print diagnostic information
  cat("Total mutations:", total_mutations, "\n")
  cat("Max generation:", max_gen, "\n")
  cat("Calculated rate:", rate, "\n")
  
  return(rate)
}
# ===============================================
# Load Required Scripts
# ===============================================
source("mutation_analysis.R")
source("C:/Users/dimet/OneDrive/Documents/DataScienceWithR/quantum_lotka_volterra.R")

# ===============================================
# Stage 1: Data Setup and Initial Processing
# ===============================================
directory_path <- "C:/Users/dimet/OneDrive/Documents/Entangled Evolution/doi_10_5061_dryad_mh206__v20190323 (1)/wgs_data/wgs_data/annotations"

# Process and save initial data
results <- analyze_mutations(directory_path)
saveRDS(results, "intermediate_results.rds")

# ===============================================
# Stage 2: Calculate Mutation Rates
# ===============================================
# Set constants
total_mutations <- 97669
max_gen <- 2907

# Calculate background rate
background_rate <- total_mutations / (max_gen * GENOME_SIZE)

# Print rate comparison
cat("Background rate:", background_rate, "\n")
cat("Drake's constant:", DRAKES_CONSTANT, "\n")

# ===============================================
# Stage 3: Statistical Analysis
# ===============================================
# Perform Fisher's tests
drake_results <- perform_fisher_test(results$raw_data, gene_lengths, DRAKES_CONSTANT)
observed_results <- perform_fisher_test(results$raw_data, gene_lengths, background_rate)

# Print sample comparison
cat("\nContingency Tables Comparison for pykF:\n")
cat("\nDrake's test:\n")
print(drake_results$pykF$contingency_table)
cat("\nObserved rate test:\n")
print(observed_results$pykF$contingency_table)

# ===============================================
# Stage 4: Analysis and Results Storage
# ===============================================
# Perform selection analysis
selection_analysis <- analyze_gene_selection(results)
saveRDS(selection_analysis, "selection_analysis.rds")

# Perform quantum Lotka-Volterra analysis
quantum_lv_results <- analyze_quantum_lv(results$raw_data)
saveRDS(quantum_lv_results, "quantum_lv_results.rds")

# ===============================================
# Stage 5: Results Output
# ===============================================
# Print Fisher test results
cat("\nFisher Test Results:\n")
cat("\nUsing Drake's constant:\n")
for (gene in names(drake_results)) {
  cat("\nGene:", gene, "\n")
  print(drake_results[[gene]])
}

cat("\nUsing observed background rate:\n")
for (gene in names(observed_results)) {
  cat("\nGene:", gene, "\n")
  print(observed_results[[gene]])
}


# Constants
DRAKES_CONSTANT <- 2.2e-10  # Mutations per site per generation
GENOME_SIZE <- 4.6e6        # E. coli genome size
TOTAL_MUTATIONS <- 97669    # Total observed mutations

# Calculate expected mutations using Drake's constant
calculate_expected_drake <- function(gene_length, generations) {
  # Expected mutations = rate * sites * generations
  expected <- DRAKES_CONSTANT * gene_length * generations
  return(expected)
}

# Calculate expected mutations using observed background rate
calculate_expected_background <- function(gene_length, total_mutations) {
  # Calculate background rate based on total mutations
  background_rate <- total_mutations / GENOME_SIZE
  expected <- background_rate * gene_length
  return(expected)
}

# Create contingency table for Fisher's test
create_contingency_table <- function(observed_gene, gene_length, generations, rate_type = "drake") {
  if (rate_type == "drake") {
    expected_gene <- calculate_expected_drake(gene_length, generations)
    # Rest should be total observed minus gene observations
    expected_background <- TOTAL_MUTATIONS - expected_gene
    observed_background <- TOTAL_MUTATIONS - observed_gene
  } else {
    expected_gene <- calculate_expected_background(gene_length, TOTAL_MUTATIONS)
    expected_background <- TOTAL_MUTATIONS - expected_gene
    observed_background <- TOTAL_MUTATIONS - observed_gene
  }
  
  contingency <- matrix(
    c(observed_gene, expected_gene,
      observed_background, expected_background),
    nrow = 2,
    byrow = TRUE
  )
  
  return(contingency)
}

# Perform Fisher's test with enrichment calculation
perform_fisher_analysis <- function(gene_name, observed_count, gene_length, generations) {
  # Drake's constant analysis
  drake_table <- create_contingency_table(
    observed_count, 
    gene_length, 
    generations, 
    "drake"
  )
  
  # Background rate analysis
  background_table <- create_contingency_table(
    observed_count,
    gene_length,
    generations,
    "background"
  )
  
  # Perform tests
  drake_test <- fisher.test(drake_table)
  background_test <- fisher.test(background_table)
  
  # Calculate enrichment scores
  drake_enrichment <- log2(observed_count / drake_table[1,2])
  background_enrichment <- log2(observed_count / background_table[1,2])
  
  return(list(
    gene = gene_name,
    drake_table = drake_table,
    background_table = background_table,
    drake_results = drake_test,
    background_results = background_test,
    enrichment = list(
      drake = drake_enrichment,
      background = background_enrichment
    )
  ))
}

# Example usage for pykF
pykF_analysis <- perform_fisher_analysis(
  gene_name = "pykF",
  observed_count = 210,
  gene_length = 1413,  # pykF length in bp
  generations = 2907   # Max generations in study
)

# Print contingency tables
print(pykF_analysis$drake_table)
print(pykF_analysis$background_table)
```
